# Quantifying Cell-type-specific Differences of Single-cell Datasets using UMAP and SHAP

**DOI:** 10.1101/2022.07.15.500285

**Authors:** Hong Seo Lim, Peng Qiu

**Affiliations:** Department of Biomedical Engineering, Georgia Institute of Technology, Atlanta, GA, USA

## Abstract

With the rapid advances in single-cell profiling technologies, larger-scale investigations that require comparisons of multiple single-cell datasets can lead to novel findings. Specifically, quantifying cell-type-specific responses to different conditions across single-cell datasets could be useful in understanding how the difference in conditions is induced at a cellular level. Here we present a computational pipeline that quantifies the cell-type-specific differences and identifies genes responsible for the differences. We quantify differences observed in a low-dimensional UMAP space as a proxy for the difference present in the high-dimensional space and use SHAP to quantify genes driving the differences. Here we applied our algorithm to the Iris flower dataset, scRNA-seq dataset, and mass cytometry dataset, and demonstrate that it can robustly quantify the cell-type-specific differences and it can also identify genes that are responsible for the differences.

## 1. INTRODUCTION

Traditional single-cell profiling technologies such as flow cytometry and mass cytometry have been widely used to understand the cellular diversity through the measurements of multiple proteins of the cells [1-3]. Rapid advances in single-cell RNA-sequencing (scRNA-seq) technologies have provided novel opportunities for new insights and discoveries by unveiling cellular heterogeneity at single-cell resolution [4-7]. Various technologies for scRNA-seq have been developed in recent years, and a great number of scRNA-seq data are now routinely generated [8].

With the growing amount of available single-cell data, it is now feasible to conduct larger-scale investigations that require combining or comparing multiple single-cell datasets, which can potentially lead to novel findings. More specifically, comparisons of various experimental conditions, such as control vs. stimulation or healthy vs. disease, can unveil new findings that would have not been possible if the analysis were focused on only one sample or one condition [9-11]. Such comparisons based on single-cell data can be used to detect cell-type-specific biological responses to various conditions and elucidate which cell types are subject to greater change due to the condition examined.

Here we present a new pipeline that could (1) quantify cell-type-specific responses across multiple scRNA-seq datasets and conditions, and (2) identify genes associated with the cell-type-specific responses. The core idea of this pipeline is to examine single-cell datasets in the UMAP [12] space, and quantify the differences in distributions of individual cell types between various conditions. For instance, if we have two datasets that are nearly identical, the two datasets would be inseparable in the UMAP visualization generated from the concatenation of the two datasets. On the other hand, if we have two datasets that harbor systematic or cell-type-specific differences in its original high-dimensional space, concatenation of the datasets followed by UMAP will show visually obvious differences in the UMAP space. Hence, we quantify differences observed in a low-dimensional UMAP space as a proxy for the difference present in the high-dimensional space. When the differences are systematic, such as the batch effect that would affect all cell types, the UMAP space will just show a clear separation of the two datasets, which is often not biologically meaningful. In contrast, when the differences are biological and cell-type-specific, the UMAP space can reveal which cell types are more affected by the conditions, because the UMAP space is able to preserve similarities among data points (cells) in the original space (Figure 1.a). After transforming the data from the original gene space to the UMAP space, a machine learning model, XGBoost, is trained to predict each cell’s UMAP coordinates based on its gene expression profile. XGBoost [13] is a state-of-the-art machine learning algorithm based on the gradient boosting ensemble method, and it has been widely used to analyze genomic data in various settings. Upon the training of the XGBoost model, Shapley Additive exPlanations (SHAP) analysis is applied to the model. SHAP is designed to explain the output of the machine learning model through a game-theory-based approach, and its usage is specifically optimized for tree-based learning models [14]. For each cell, SHAP analysis can quantify each gene’s contribution to the coordinates where that cell is projected in the UMAP space. The SHAP values of cells in a specific cell type can be used to quantify which genes are associated with the changes of this cell type in response to the conditions being compared. A flowchart of our cell-type-specific variation quantification pipeline is described in Figure 1.b. Using several sets of data in various experimental contexts, we demonstrated that the proposed pipeline could robustly quantify cell-type-specific differences and identify cell-type-specific genes associated with the variations.

**Figure 1.**
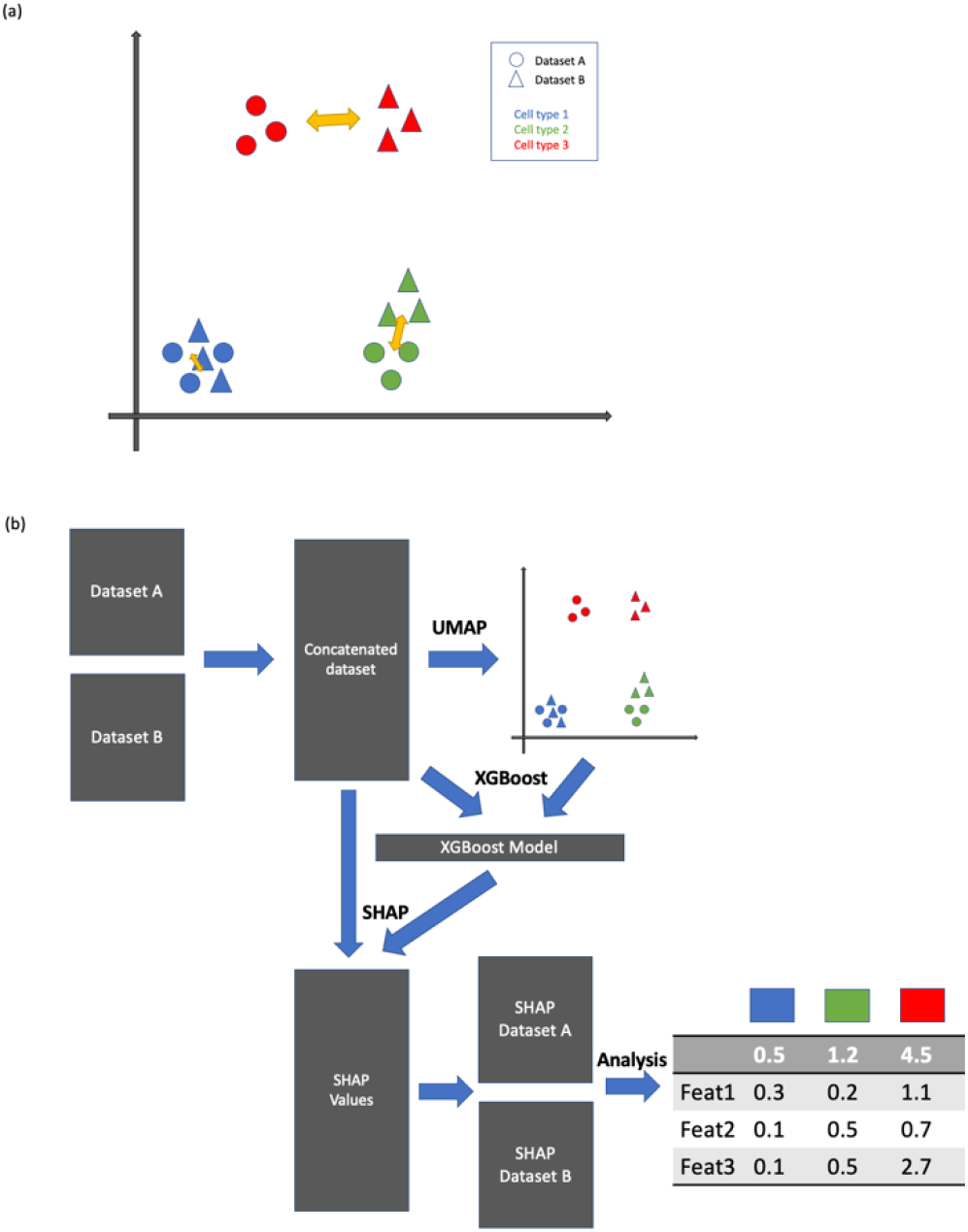
Overview of suggested pipeline. (a) Demonstration of difference of certain label in low-dimensional space. (b) flowchart of the suggested pipeline.

## 2. METHODS

In this chapter, the following method is tailored for the comparisons of scRNA-seq datasets, yet the usage of the suggested pipeline is not limited to scRNA-seq. Any sets of datasets generated by the same technology (for instance, flow cytometry vs. flow cytometry or mass cytometry vs. mass cytometry), that share certain features that can be used for the concatenation of the set of datasets followed by UMAP visualization can be applied.

### 2.1 Data preprocessing

Our pipeline requires two or more datasets to compute cell-type-specific differences across the datasets. For each scRNA-seq dataset, we perform initial filtering of cells (min_genes = 2000) and filtering of genes (min_cells=3). Cells with high mitochondrial expression (5 percent) or too many total counts (n=2500) are also removed. The filtered dataset undergoes library size normalization (target_sum=10,000), log transformation, and identification of highly variable genes. Since each dataset has its own unique highly variable gene list, we compute the union of the highly variable gene lists, and use the union in subsequent steps of our pipeline.

### 2.2 Concatenation of datasets and dimensionality reduction

Given the union of highly variable genes in multiple datasets under consideration, we filter each dataset and reshuffle the rows (genes) so that each dataset contains single-cell expression data of the union of highly variable genes, and the order of the genes is the same for each dataset. The filtered datasets are concatenated to be one dataset. Dimensionality reduction is then applied to this concatenated dataset. More specifically, principal component analysis (PCA) followed by UMAP is applied to the dataset. The resulting UMAP coordinates go through a normalization step where we scale each axis from 0 to 10. This normalization step ensures that every run of our pipeline has comparable measurements in the UMAP space. The result of the dimensionality step is the two-dimensional UMAP coordinates of all cells across all datasets under consideration. The coordinates serve as an input to the subsequent SHAP analysis.

### 2.3 XGBoost to learn UMAP coordinates and gene expression

We concatenate the two datasets using the normalized and initially filtered datasets (from step 2.1). The combined dataset is an input to the XGBoost machine learning model. The model is trained to predict each cell’s UMAP coordinate (from step 2.2) using the gene expression data. Since the goal of the XGBoost model is to learn the relationship between the UMAP coordinates and gene expression within the dataset, and the model is not intended to be used to make predictions for other datasets, the model is trained with the entire dataset. Upon the completion of the fitting of the model, the model becomes another input to the subsequent SHAP analysis.

### 2.4 SHAP analysis

SHAP analysis requires two inputs: model and data. SHAP analysis generates SHAP values from the inputs, and the dimension of the SHAP values is the same as the input data. For each cell, its SHAP values quantify each gene’s contribution to determine its coordinate in the UMAP space. In our pipeline, the trained XGBoost model and the concatenated datasets (from step 2.3) are the inputs to the SHAP analysis.

### 2.5 Quantifying cell-type specific difference from SHAP values

Because SHAP values generated from SHAP analysis have the same size (number of cells by number of genes) as the input dataset, we can filter the SHAP values by datasets and by cell type label. For each cell type label shared between two datasets, we can find two sets of SHAP values whose corresponding cells match the cell type label in respective datasets. For each set of SHAP values corresponding to the cell type label, we calculate the mean SHAP values per gene across the cells, resulting in a vector of a length equal to the number of genes, which quantifies in average how much each gene contributes to the UMAP location of the cell type in one dataset. With two datasets, we have two SHAP vectors for the specific cell type label, and we quantify the difference by taking the absolute element-wise difference between the two vectors and computing the sum of the element-wise difference. This serves as a score to quantify the difference in terms of one specific cell type across two datasets. The above steps are repeated for every cell type label, so that each label receives such a score.

### 2.6 Random permutation and correction of the scores

To evaluate the statistical significance of the scores calculated from the previous step, the same method is applied to randomly shuffled SHAP values. The random shuffling of dataset membership and the subsequent variance scoring is repeated 1000 times. The score without the shuffling is compared with the distribution of the 1000 scores from shuffled data, so that we can use the 1000 scores from shuffled data to form a null distribution to derive a p-value for the observed score without shuffling. With the random permutations, we can provide a corrected version of the original scores by subtracting the mean of the random shuffling from the original score.

### 2.7 Rank-ordering genes for each cell type

The above score quantifies the cell-type-specific difference between datasets for a specific cell type label. It is often informative to examine the top genes that contribute most to the cell-type-specific differences. Hence, we rank-ordered the absolute difference between the two SHAP vectors and show each gene’s individual score which could serve as valuable information to understand the biological interpretation of the difference detected.

### 2.8 Datasets

In this study, we used 3 different datasets to demonstrate the utility of our pipeline for quantifying cell-type-specific differences. The first dataset—Iris Dataset— is used as proof of concept. The second dataset is a scRNA-seq PBMC IFN-B stimulation data[11]. And third datasets are mass cytometry dataset of human hematopoiesis in different perturbation conditions [10].

## 3. RESULTS

### 3.1 Demonstration of cell-type specific difference quantification on the Iris dataset

The Iris flower dataset is a multivariate dataset (150 samples by 4 features) composed of 50 samples from each of three different types of iris flowers. The dataset contains four features: petal length, petal width, sepal length, and sepal width. To demonstrate our suggested pipeline, we simulated two scenarios. (1) In the first scenario (noise level = 0), the 150 samples are randomly divided into two independent datasets. (2) In the second scenario (noise level = 1), the 150 samples are randomly divided into two datasets, and then gaussian noise is added to one of the datasets. For each flower label type, our pipeline provides three pieces of information. As shown in the right panel of Figure 2, the first row shows the actual scores based on the difference in SHAP values for each label. Each dot in the plot is a feature, and its coordinate is based on the average SHAP values from each dataset. If two datasets are similar, the dots of features are likely to align well along the diagonal line, and if two datasets are different, the dots of features could deviate from the diagonal line, and the extent of deviation could indicate the contribution of certain features to the difference observed. The second row in the right panel of Figure 2 is the comparison (p-value) of the score with the scores of random shuffling shown in the histogram. The dotted line on the histogram marks the score and having a p-value that is not significant (for instance, p-value > 0.05) might suggest that the scores are statistically insignificant compared to random shuffling. The third row in the right panel of Figure 2 shows the random-noise corrected scores where the scores are subtracted by respective means from the random shuffling. For two different scenarios mentioned above, the UMAP visualizations of the datasets are shown in the left panels of Figure 2.a. and Figure 2.b.

**Figure 2.**
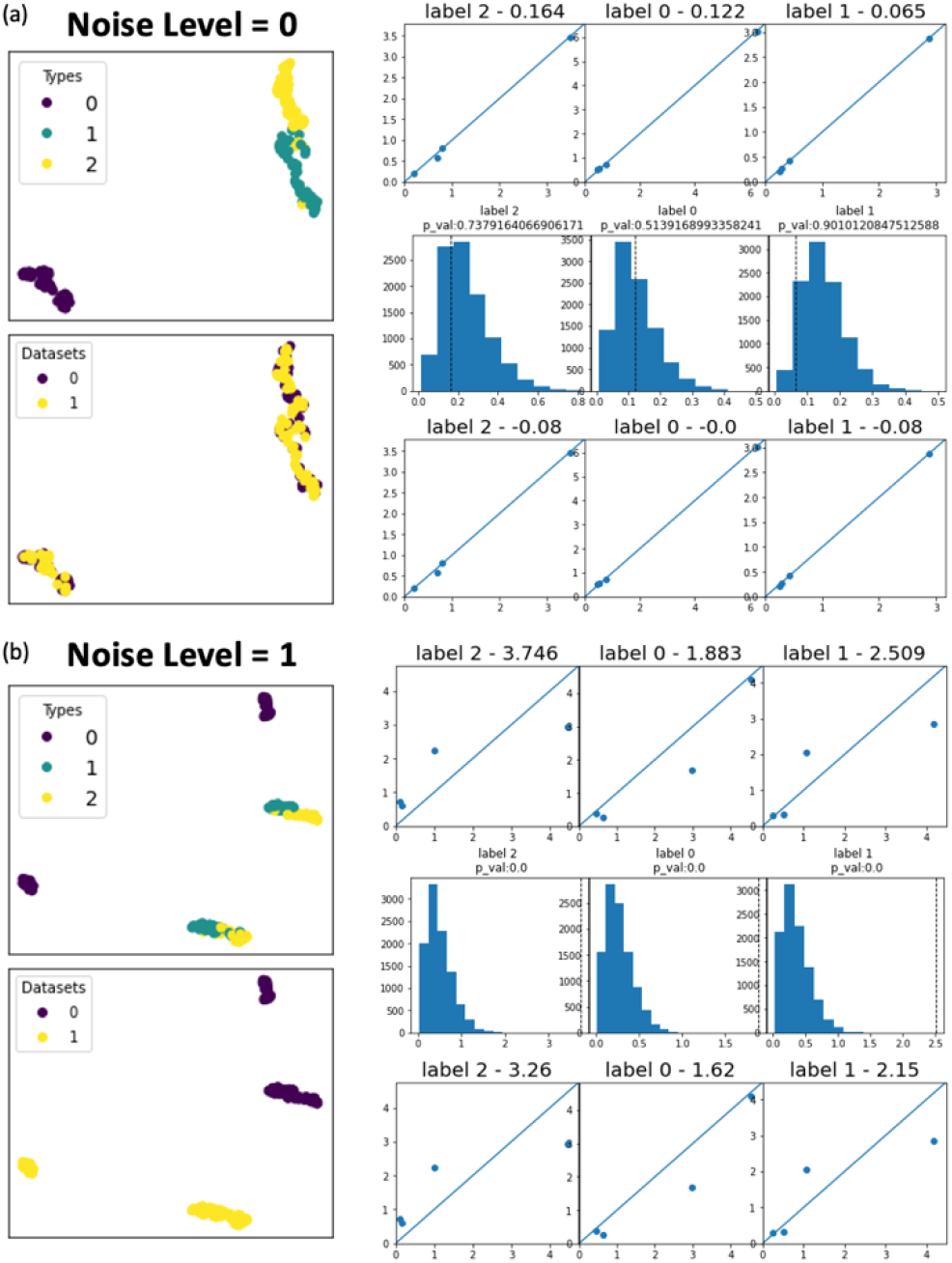
Results of the simulated IRIS datasets (a) outputs of the pipeline when added noise is 0. (b) outputs of the pipeline when added noise is 1.

For the first scenario, since the datasets are divided randomly without any perturbation, the two datasets are visually inseparable in the UMAP space (Figure 2.a). On the contrary, in the second scenario, because of the added noise on one of the datasets, two datasets are easily separable in the UMAP space (Figure 2.b). Subsequent scores of different flower types from the first scenario are close to 0 (first row of 2.a) with all dots representing features aligning closely with the diagonal line. The scores are within the range of scores from random shuffling of the SHAP scores (second row of 2.a) reflected by high p-values. The scores after random-noise corrections are even closer to zero (third row of Figure 2.a). Such results suggest that the two datasets in the first scenario are not showing meaningful differences between the two datasets which is a correct interpretation given that the two datasets were randomly split without any systematic perturbation. On the other hand, results from the second scenario show much higher scores for all flower types, and the higher scores are originated from the dots of features deviating from the diagonal lines in the scatter plots (first row of Figure 2.b). Comparisons with random shuffling scores suggest a significant difference in the datasets, illustrated by extremely small p-values (second row of Figure 2.b). Even after the random noise effects are corrected, the scores are still significantly greater than the scores from the first scenario (third row of Figure 2.b). Given we added systematic noise to one of the datasets in the second scenario, the results, and the interpretations correctly capture the difference simulated in these datasets.

To demonstrate the power of our pipeline, we prepared more specific perturbation scenarios using the same dataset. In the first scenario, we randomly split the original data into two, and we added Gaussian noise to only one feature in one of the datasets. The results for this feature-specific perturbation are shown in Figure 3.a. Here the perturbed feature was petal width, and the perturbance resulted in a visual difference between the two datasets in the UMAP space. Random-noise corrected scores (second row of Figure 3.a) suggest some significant difference between the datasets even after random-noise correction, and we noticed that a specific flower type (i.e. Label 0) was more affected by the perturbance of the feature. Lastly, the table in Figure 3.a summarizes each feature’s contribution for each label type, and it shows that the petal width is in fact the most contributing (to the difference) feature across all flower types, hence correctly capturing the perturbation we simulated.

**Figure 3.**
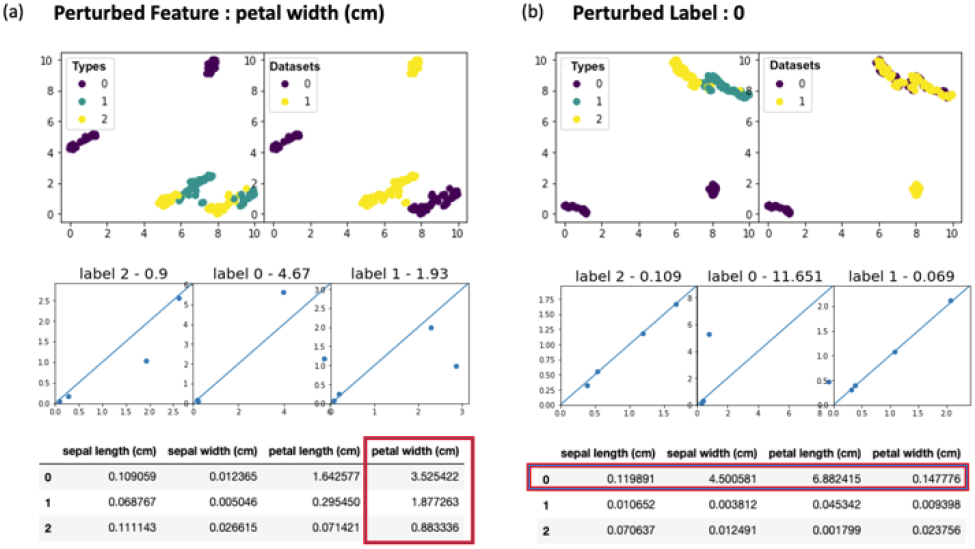
Feature-specific and label-specific perturbation. (a) UMAP visualization(top), label scores and scatter plots(middle), and feature scores(bottom) for the perturbed feature scenario. (b) UMAP visualization(top), label scores and scatter plots(middle), and feature scores(bottom) for the perturbed label scenario.

In the second scenario, we randomly split the original data into two, and we added Gaussian noise to all features of one flower type only. The results of the label-specific perturbation of flower type 0 are described in Figure 3.b. The UMAP visualization illustrates that only label 0 is visually differentiable among the labels as shown in Figure 3.b. The subsequent analysis demonstrates a significantly higher score for label 0, and large deviations of the features from the diagonal line in the scatter plots were also observed for label 0. The table in Figure 3.b shows that all features of label 0 are orders of magnitude greater than other labels, hence again correctly capturing the very perturbation we simulated.

### 3.2 IFN-B Stimulation of Human PBMC

We applied our pipeline to human PBMC samples under two different conditions: one sample stimulated with interferon-beta (IFN-B) and another sample unstimulated serving as a control [11]. When concatenated after the preprocessing step, the two datasets exbibit a large batch difference as shown by the completely separate locations in the UMAP visualizations as shown in Figure 4.a. Using the provided cell type labels, we observed that the difference is not specific to certain cell types, but all cell types were collectively affected as shown in Figure 4.a.

**Figure 4.**
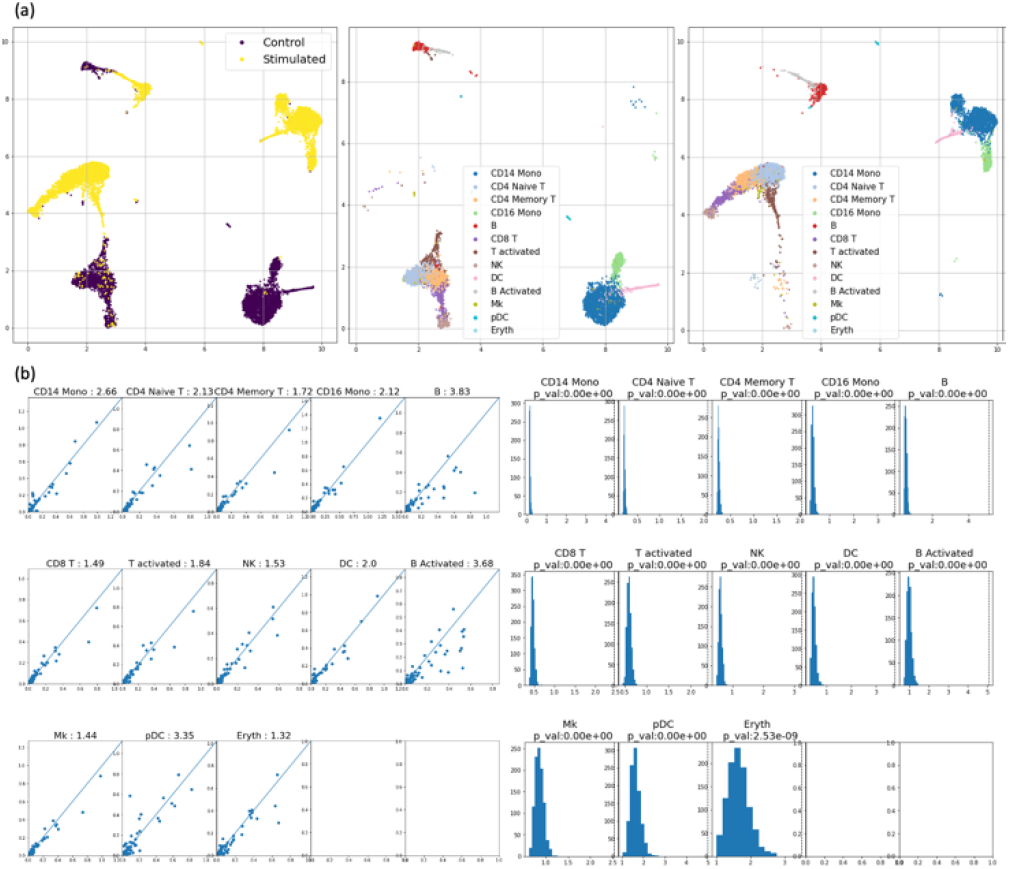
PBMC stimulated vs. unstimulated. (a) UMAP visualizations of the two datasets. (b) Noise-corrected scores and scatter plots(left), and histograms of the shuffled SHAP values.

Our subsequent analysis quantified such differences shown in the UMAP. In Figure 4.b, we showed scatter plots and noise-corrected scores (left) and comparisons with scores of random shuffling. All cell types received high variation scores (>1.3), and the comparisons with randomly shuffled scores suggest that all scores are statistically significant signals. Such universal changes by interferon-beta stimulation were also seen in the previous studies whose analysis were done on the same set of datasets [11, 15].

To further demonstrate the utility of our gene scores in reflecting each gene’s contribution to the difference in the datasets, we took a closer look at some marker genes of human PBMC. First, we used SHAP difference scores for cell-type-specific marker genes, and our scores suggest that the marker genes for specific cell types received higher SHAP scores as shown by the the cell type by gene matrix in Figure 5.a. In addition, we checked the scores for IFN-B genes (ISG15, IFI6, IFIT1, IRF7, MX1, OAS1) that are known to be affected uniformly across all cell types.

**Figure 5.**
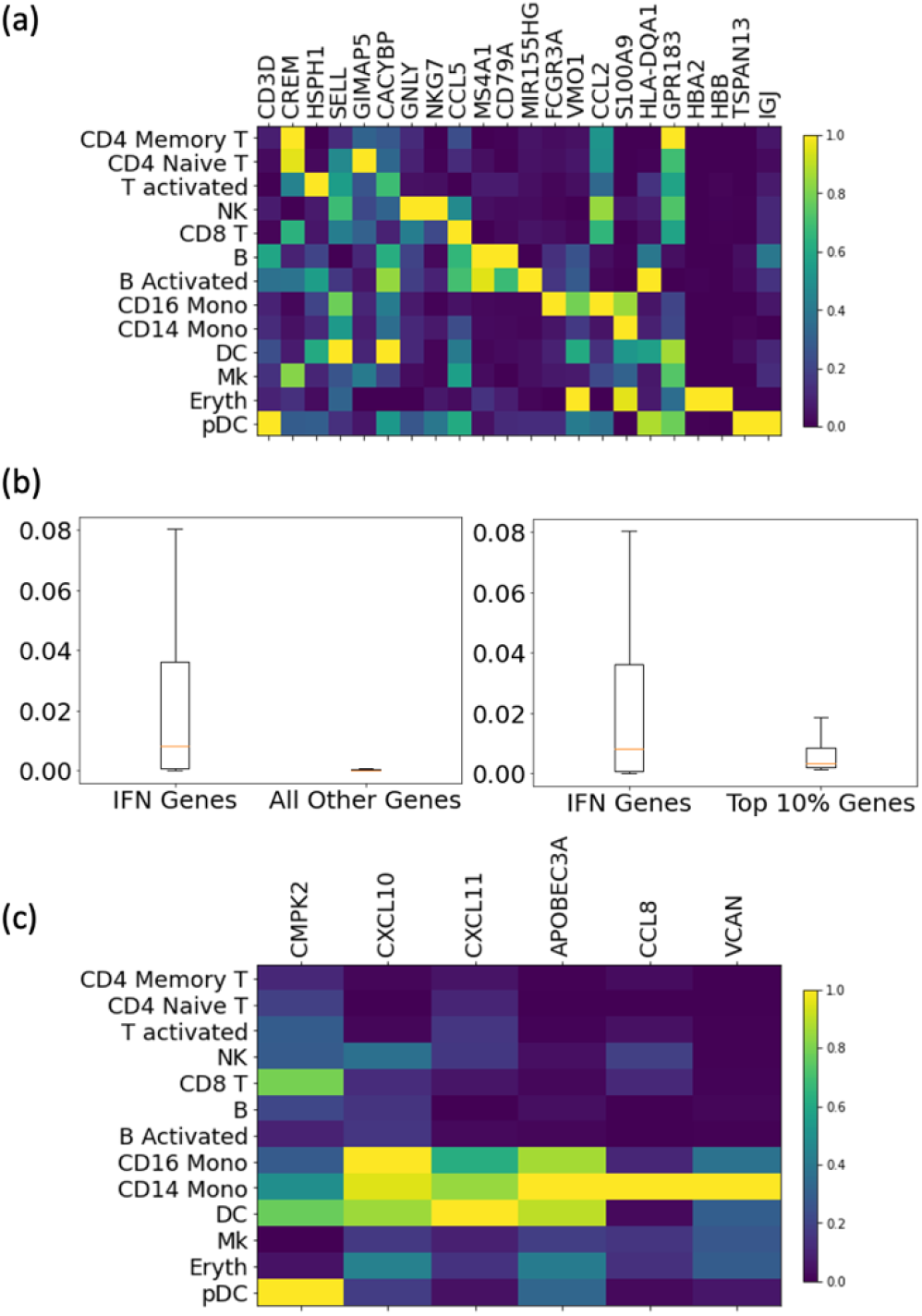
Individual gene scores for stimulated vs. unstimulated PBMC. (a) Individual gene scores for cell-type marker genes. (b) Distribution of gene scores between IFN genes vs other genes. (c) Gene scores for cell-specific IFN genes across the cell types.

Comparing scores from this list of genes across all cell types with the scores for all other genes, we observed that scores from the IFN-B genes were significantly elevated compared to other genes as shown in Figure 5.b. Furthermore, we compared the IFN-B gene scores with the top 10% of genes with highest scores, and the scores of IFN-B genes are still comparable to the genes received high scores, suggesting the elevated contribution of the IFN-B genes to the difference we observed in the two datasets. Lastly, we checked the individual scores for cell-type-specific IFN-B marker genes (CMPK2, CXCL10,CXCL11, APOBEC3A,CCL8,VCAN). The result is shown in Figure 5.c, where certain cell types show greater responses to certain genes. For example, CD16 monocytes and CD14 monocytes were affected more by CXCL10, CXCL11, CCL8, and VCAN. Such findings were also described in the previous study [15], demonstrating the utility of our pipeline.

### 3.3 Mass cytometry of human hematopoiesis under different perturbation

Next, we applied our pipeline to mass cytometry of human hematopoiesis datasets of bone marrow acquired from Bendall et. al [10]. Specifically, we used 4 unstimulated samples, one sample under GM-CSF (granulocyte/macrophage colony-stimulating factor) stimulation, and one sample under TNFa (tumor necrosis factor alpha) stimulation. Datasets contain cell type label information from the previous study. For each stimulated sample, we conducted four independent runs of our pipeline, one for each unstimulated sample and the stimulated sample. Since each run would generate a difference score for each cell type label, quantifying the changes of one cell type in response to a stimulation, we summarized the distribution of the scores per each cell type label across the four comparisons as shown in the left panel of Figure 6.a. Monocyte, myelocyte, pDC, pro-myelocytes, early monocyte, and megakaryocyte were cell types that had highest changes due to the GM-CSF stimulation, and the findings are congruous with earlier studies highlighting activation of myeloid cells under GM-CSF stimulation [16]. Furthermore, we focused on the contribution of cell functional markers in the difference scores we observed, and STAT5 received high contributing scores among the highly scored cells as shown in the left panel of Figure 6.b. Previous studies have demonstrated that GM-CSF signals through STAT5, hence our findings reaffirm the previous findings [17, 18], and it demonstrates that our pipeline not only scores difference per cell-types but also identifies markers driving the difference.

**Figure 6.**
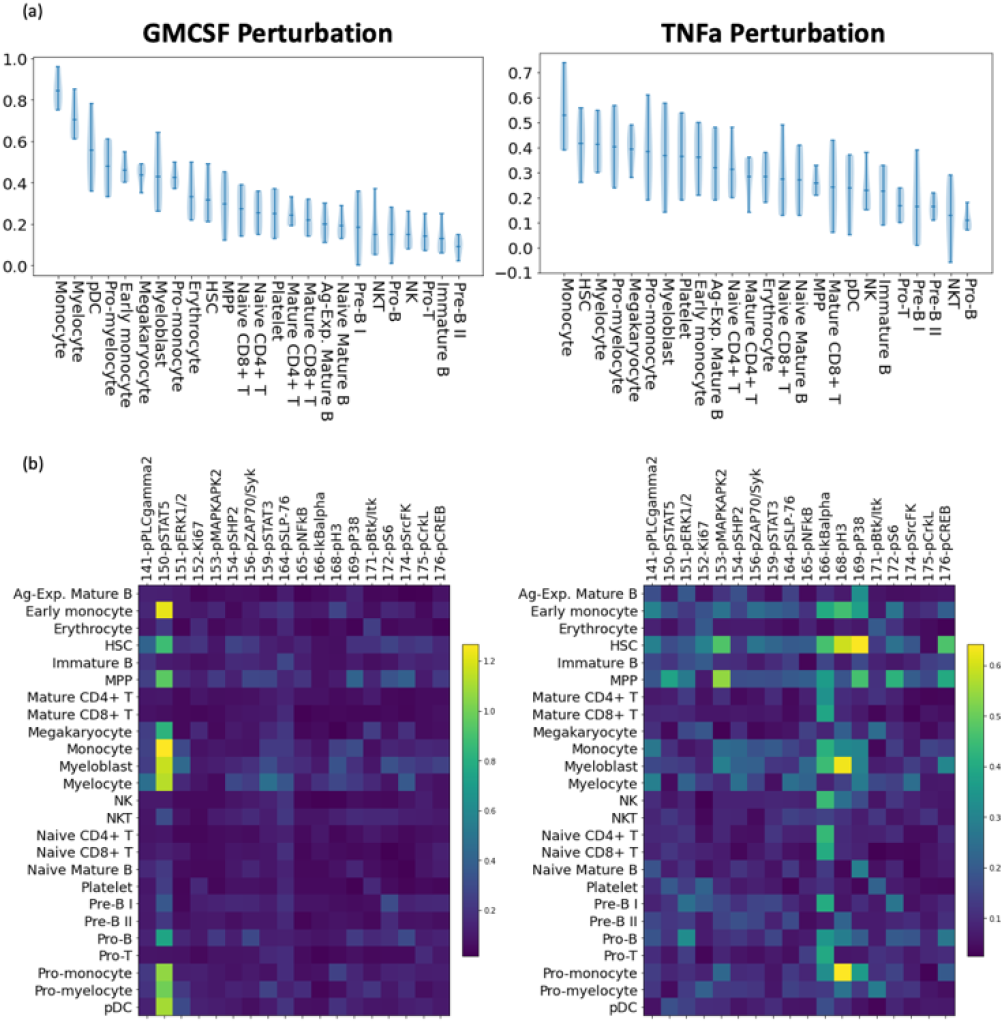
Human bone marrow hematopoiesis results. (a) violin plots of cell type’s scores from comparisons with unstimulated samples with the stimulated sample. (b) SHAP difference scores for cell functional markers across the cell types.

The same analysis was applied to unstimulated samples and a sample with TNFa stimulation. As seen in the right panel of Figure 6.a, monocyte, HSC, myelocyte, and pro-myelocytes were among the top cell types that showed high difference scores, and the findings are again congruous with myeloid-derived changes observed in TNF signaling [19]. The contribution of cell functional markers as shown in Figure 6.b suggest that IKBalpha is the one marker contributing the most to the difference we observed across many myeloid-lineage cell types. IKBalpha is a protein that inhibits NFKB transcription factor, where NFKB is a transcription regulator known to be directly activated by TNFa cytokines, hence confirming our finding [20].

## 4. DISCUSSION

Here, we present a new pipeline to quantify cell-type-specific differences and identify genes driving the difference. Our pipeline exploits the quantifiable differences seen in the low-dimensional UMAP and used SHAP analysis to measure the difference. We have demonstrated that our algorithm could correctly capture various perturbation scenarios—systematic variation across all cells, a variation on a specific feature, a variation on a specific label—as seen from the simulated datasets based on the Iris flower dataset. Then we showed that our algorithm correctly captures biological responses to stimulation in human hematopoiesis bone marrow mass cytometry datasets. We demonstrated the algorithm’s utility in interpreting and quantifying differences in various scRNA-seq datasets where our results agree with previous studies. We believe that our suggested pipeline is intuitive in the sense that we are quantifying the difference seen in UMAP visualization and quantifying how each gene contributes to the difference. Because our analysis is based on the difference we can visually see, it makes results from our analysis readily interpretable.

There are several limitations of this work. First, since our analysis is dependent on robust and accurate UMAP projections. If certain differences in high-dimensional space were not reflected in the UMAP projections, our analysis would not be able to capture them. Second, individual SHAP contribution scores could be affected by the curse of dimensionality in scRNA-seq data. The individual SHAP scores of genes are dependent on what XGBoost learns from the relationship between UMAP coordinates and gene expression profiles, and if XGBoost fails to utilize all correct features due to many features, and the existence of multicollinearity, some features could be potentially not identified by our pipeline

As for the future works, our pipeline can be improved by applying additional dimensionality reduction runs (UMAP with different parameters or tSNE) to enhance robustness, and by regulating XGBoost to use a series of a subset of genes when training to handle the so that important yet potentially left-out genes could be correctly used in the training.

Overall, our pipeline provides a thorough statistical approach to quantifying cell-type-specific differences. While most computational tools aim to correct the differences among the datasets, our pipeline embraces the differences and exploits the differences.

